# MAGE enables population level RNAseq driven genotyping and (differential) allelic divergence detection in healthy kidney and carcinoma

**DOI:** 10.1101/2022.09.06.506720

**Authors:** Stroobandt Cedric, Goovaerts Tine, Coussement Louis, De Graeve Femke, Voorthuijzen Floris, Van Steenbergen Laure, Galle Jeroen, Van Criekinge Wim, De Meyer Tim

**Author notes:** Corresponding author Tim De Meyer. Shared first authors (Stroobandt Cedric, Goovaerts Tine).

## Abstract

Decreasing sequencing costs have instigated large-scale RNAseq experiments, yet genetic polymorphisms in such data remain poorly exploited. Currently, allele-specific expression (ASE) studies focus almost exclusively on genetic variants explaining expression differences (*cis*-eQTLs), largely ignoring other ASE effects. The latter are typically associated with higher variance in expression of both copies of a gene, here called Allelic Divergence (AD). We therefore developed an RNAseq-driven population-level beta-binomial mixture model for (differential) AD detection. The model simultaneously enables RNAseq-driven genotyping, which outperforms alternative RNA genotyping methods when applied on healthy kidney data from The Cancer Genome Atlas. Moreover, we identify well-known non-*cis*-eQTL ASE, e.g. random monoallelic expression of HLA and immunoglobulin genes in healthy kidney, as well as allele-specific aberrations in clear cell kidney carcinoma, including long-range epigenetic silencing of protocadherins, copy-number alterations, and loss of imprinting. These methods are available as the Modeller of Allelic Gene Expression (MAGE) tool suite: https://biobix.github.io/MAGE/.

## 1. Introduction

RNAseq is typically used in differential gene expression studies, relying on well-established methods such as *EdgeR*^1^ and *limma-voom*^2^. Yet, similar to DNAseq, it also captures genetic variation – e.g. single nucleotide polymorphisms (SNPs)^3^ – and thus allows for differentiation between alleles at polymorphic loci^4^. This enables the study of genes for which alleles feature unequal expression, i.e. allele-specific expression (ASE)^5,6^, with high-impact applications in e.g. plant genetics^7^, autoimmunity^8^, cancer^9^ and evolutionary studies^10^. ASE can be attributed to genetic variation of alleles themselves, consistently manifesting as unequal expression between allelic variants, here termed Allelic Bias (AB). This is the case for *cis* expression quantitative trait loci (*cis*-eQTLs)^4^, though AB can also be caused by technical phenomena like alignment bias. In contrast, other ASE effects are independent of the underlying genetics and thus, while they still introduce unequal expression of both copies of a gene within an individual, they don’t manifest as unequal expression between allelic variants in a population but rather as an additional source of variance. We denote this specific class of variance-acting ASE phenomena as “Allelic Divergence” (AD). AD then includes genomic imprinting, in which only the allele of one specific parent is expressed. Imprinted genes typically control growth and development and their dysregulation has been demonstrated in both congenital and acquired diseases, including cancer^11^. For AD loci featuring random monoallelic expression (RME), e.g. X-chromosome inactivation, the expressed allele is selected at random rather than in a parent-of-origin-specific manner^12^. Finally, also disease-associated phenomena such as copy-number alterations (CNA), (promoter) mutations and epigenetic artifacts can act on separate copies of a gene. When recurring in diseased populations, this type of AD is compatible with the gene’s potential causal role in disease (progression)^13^.

Despite AD’s biological relevance, it can be challenging to detect in bulk RNA samples, as disease associated AD is often diluted by admixture of healthy cells; whereas for RME, cells expressing opposite alleles may be mixed. Nevertheless, as the effect of (ad)mixture is typically limited and given a large enough population, increased variance due to AD can be statistically detected.

Indeed, a plethora of phenomena makes comprehensive ASE studies non-trivial. Early methodologies often simplified ASE to individual-level deviations from a 1:1 allelic expression ratio^14^ at heterozygous loci, a strategy prone to technical aberrations, e.g. the aforementioned alignment bias. Population-level ASE modelling is more appropriate, yet available methods typically focus on eQTL detection rather than AD, e.g. WASP^15^. Indeed, methodologies to study AD in bulk RNAseq data are particularly rare, with disease-associated AD usually being studied on diseased datasets alone (ignoring potential “healthy background” AD) or – if including controls - being limited to paired data^13^. Besides these limitations, a final drawback of most ASE-modelling tools is the reliance on genotyping data, be it via (phased) DNAseq or SNP arrays, often even for parent-offspring trios, to identify heterozygous individuals. A sufficiently large number of the latter is required to study ASE per locus, which implies large population sizes, especially in human populations, where breeding designs to increase heterozygosity are infeasible. The reliance on genotyping data is therefore an expensive one, doubly so as it reduces data-usage efficiency by only being able to study RNAseq data covered by the genotyping assay^16^. In summary, to date, no population-level methods enable modelling of multiple ASE-effects, let alone one solely relying on RNAseq for cost- and data-efficiency.

We previously modelled monoallelic gene expression using solely population-level RNAseq data to study loss of imprinting in (breast) cancer^16^. As non-imprinting AD rarely leads to complete monoallelic expression, here we introduce a methodology to generically assess (differential) AD, while simultaneously developing an ASE-aware RNAseq-driven genotyper in the process. These methodologies, including an updated imprinting analysis, are integrated in the Modeller of Allelic Gene Expression (MAGE) software suite. We apply MAGE on healthy kidney tissue and clear cell renal carcinoma (KIRC) RNAseq data from The Cancer Genome Atlas (TCGA), demonstrating its genotyping and AD-screening functionalities. By evaluating available TCGA Infinium HumanMethylation 450k and CNA data, we confirm that MAGE detects all common KIRC copy-number alterations, but also monoallelic expression of the protocadherin gene clusters associated with long-range epigenetic silencing. This sheds further light on renal cell carcinoma biology, the sixth and tenth most common cancer in men and women, respectively^38^.

## 2. Results

Section 2.1 introduces how RNAseq allele counts can be modelled as a mixture of beta-binomial distributions to enable DNA-free genotyping and (differential) AD studies, complemented with an update of our previously developed (loss of) imprinting methodology^16^. In Section 2.2, we illustrate the performance on renal cancer RNAseq data provided by TCGA, using corresponding SNP array, CNA as well as Infinium HumanMethylation 450k data for benchmarking.

### 2.1. MAGE’s statistical models

As input data, MAGE starts from per-sample reference- and variant-SNP counts for each locus-of-interest. Note that MAGE is currently limited to modelling SNP-level data with two variants, here referred to as alleles.

MAGE’s analyses can be divided into four categories, illustrated in Figure 1 using histograms of per-sample reference allele fractions (reference allele count over total allele count) of a hypothetical locus-of-interest:

**Figure 1:**
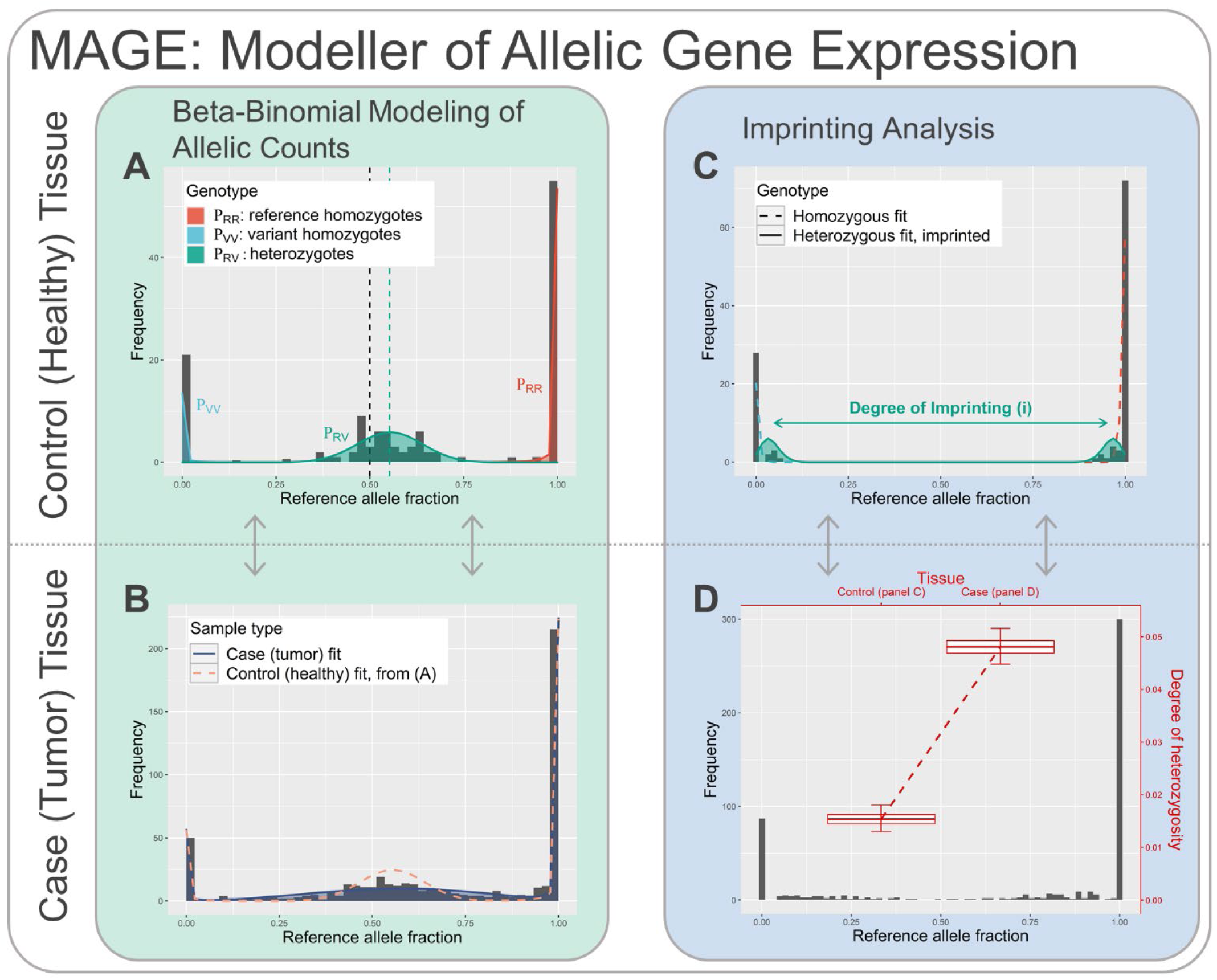
histograms of per-sample reference allele fractions of a particular locus, illustrating the different MAGE analyses. Histograms represent the observed data, lines/boxplots the fitted models.

A. ASE-aware genotyping
B. Detection of (differential) allelic divergence ((d)AD)
C. Detection of imprinting
D. Detection of loss of imprinting (LOI)

The following Sections elaborate the rationale behind the underlying models, whereas the Methods section covers in-depth mathematical- and implementation details.

### 2.1.A. ASE-aware genotyping

MAGE models each locus’ reference and variant allele counts as a mixture of beta-binomial observations, which are themselves characterized by three parameters: (1) the observed total allele count *n*, (2) the expected fraction of reference allele counts π, (3) the overdispersion parameter *ρ* capturing biological variance on top of technical variance captured by the regular binomial distribution. The total probability mass function is thus:

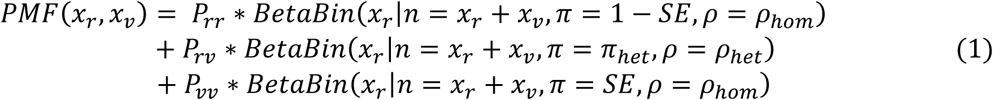

With (*x*_*r*_, *x*_*υ*_) the observed reference and variant allele counts, (*P*_*rr*_, *P*_*υυ*_, *P*_*rυ*_) relative genotype frequencies of reference- and variant-homozygotes and heterozygotes, respectively, *SE* the sequencing error rate, π_*het*_ the expected reference allele fraction in heterozygous samples, and (*ρ*_*hom*_, *ρ*_*het*_) the overdispersion of homozygous and heterozygous samples, respectively (ranging from zero to one; assumed equal for both homozygous fractions). Following our definitions from Section 1, π_*het*_ captures Allelic Bias (AB) and *ρ*_*het*_ captures the amount of Allelic Divergence (AD) present in a population’s expression data. With the exception of *SE*, which is a constant hyperparameter to be set or estimated separately in advance, MAGE fits all distributional parameters on a per-locus basis using a robust expectation-maximization (EM) algorithm (Figure 1A; vertical dashed lines denote AB).

During the fitting process, EM inherently calculates each sample’s relative probabilities to originate from each genotype’s component beta-binomial distribution in Equation 1, directly leading to RNAseq-driven genotyping. Capturing both AB and AD makes for a very flexible ASE-aware genotyping tool. Note that MAGE allows testing for significant AB (different from the balanced π_*het*_ = 0.5 scenario) using a likelihood ratio test (LRT). Yet, interpretation is complicated by the fact that AB may be explained by both technical (e.g. alignment bias) and biological (e.g. *cis*-eQTL loci) root causes.

#### 2.1.B. Detection of (differential) allelic divergence

In the introduction, we hypothesized that genes for which dysregulation is causal in a disease etiology or its progression will often feature additional AD in affected tissue as compared to healthy tissue, due to e.g. CNA, (promoter) mutations, aberrant hypermethylation, or a mix thereof. Such effects cause a shift in expressed alleles towards either (relatively) more reference or more variant reads on a per-sample basis *but* are equally likely to affect either allele at a population scale, causing increased AD to be captured by *ρ*_*het*_. Loci featuring consistent allelic dysregulation in disease can thus be identified as those loci featuring *differential Allelic Divergence* (dAD), with typically larger *ρ*_*het*_ in case compared to control tissue. Assuming that cases and controls share their remaining distributional parameters (π_*het*_, *ρ*_*hom*_, *P*_*rr*_, *P*_*υυ*_, *P*_*rυ*_; Equation 1), dAD can be detected using an LRT. This is illustrated in Figure 1B, with the dashed line representing the control sample model (*ρ*_*het*_ from panel A) and the solid line representing cases with increased *ρ*_*het*_.

#### 2.1.C. Detection of (loss of) imprinting

MAGE’s (loss of) imprinting analyses are built upon earlier work by our research group^16^, albeit with some refinements (see Section 4.1.D). The methodology focuses on the telltale characteristic of imprinting, i.e. monoallelic expression, as imprinting manifests itself as either the paternal or the maternal allele being consistently (and typically completely) silenced across samples. These methods higher cannot adequately study such extreme AD, as heterozygous samples become practically indistinguishable from homozygous ones. Instead, after estimating allele frequencies from raw allele counts, MAGE relies on population genetics to construct a Hardy-Weinberg-Equilibrium (HWE) conform (regular) binomial mixture model, in the current version adjusted for the estimated inbreeding coefficient (see Methods). In this model, the heterozygous distribution is split up according to degree-of-imprinting *i*. An *i* equal to 0 corresponds to no imprinting and a single peak of heterozygous samples whereas an *i* of 1 corresponds to complete imprinting with two heterozygous peaks undistinguishable from the homozygous ones (Figure 1C). For statistical testing, the estimated *î* can be compared against the null hypothesis (*i*=0) by means of a LRT. In practice this procedure leads to *candidate* imprinted genes as genes that show extreme RME throughout comprised cell types will also be detected. To disambiguate, trio data is required.

For (candidate) imprinted loci, MAGE identifies loss of imprinting (LOI) as those loci featuring re-expression of the silenced allele in case vs. control samples. In the current implementation, this is detected by considering the least (most) expressed allele per sample as success (failure), and performing binomial logistic regression to compare the degree of success (i.e. reexpression) between both populations (Figure 1D; Section 4.1.D). Contrasting the previous procedure, this strategy takes variable sequencing depth into account. In addition, genes displaying LOI in disease are expected to be significantly upregulated in cases compared to controls, which we previously termed canonical LOI^16^.

### 2.2. ASE case study on TCGA data

In this Section, we demonstrate MAGE’s added value by application on TCGA renal RNAseq data. More specifically, for genotyping and imprinting detection (2.1.A and 2.1.C), we use data of 128 healthy kidney samples, comprised of the control samples of Kidney Renal Clear Cell Carcinoma (KIRC; N=72), Kidney Renal Papillary Cell Carcinoma (KIRP; N=32) and Chromophobe Renal Cell Carcinoma (KICH; N=24). To avoid introducing bias through batch effects, for differential imprinting (DI) and AD studies (2.1.B and 2.1.C), only the 72 KIRC control samples are retained, and compared against TCGA’s 268 (deduplicated) KIRC stage 1 tumour samples.

#### 2.2.A. Genotyping renal RNAseq data

TCGA provides Affymetrix Human SNP Array 6.0 data (targeting 906,600 SNPs) for 126 of its 128 healthy renal samples, providing the gold standard genotypes to assess MAGE’s genotyping performance. For benchmarking, we genotyped the same RNAseq data using the standard GATK RNAseq short variant discovery pipeline^17^, as well as SeqEM^18^ which is similar to MAGE in that it relies on EM, yet uses regular binomial models and ignores AB.

Of the 68,918,146 sample-locus combinations genotyped by MAGE (only loci covered by at least 10 samples) on TCGA KIRC data, 4,021,426 are retained for an unbiased and high-fidelity comparison (Section 4.2.D); Figure 2 depicts the genotyping error rate of the three methods as a function of the applied per-SNP per-sample minimal read count filter, given the latter’s straightforward impact on genotyping accuracy. For a read count filter of 10 (1,327,562 of 4,021,426 to-be-genotyped sample-locus combinations remaining), MAGE’s genotyping error rate (1.24%) is 43% lower than GATK’s (2.17%) while that of SeqEM lies in between (1.46%). MAGE’s relative performance is especially remarkable when including lower-count data (count filter 3-9) while maintaining its advantage, especially over GATK, on higher-count data as well. Of course, MAGE’s reliability can be improved via model-fit-dependent filters, the most important one being based on (inbreeding coefficient-adjusted) HWE (see Methods), as loci showing strong HWE-deviation are more likely to feature poor quality or interfering biological effects (e.g. imprinting). When combined with a read count filter of 10, the HWE-filter leads to further genotyping error rate improvement (1.24% to 1.16%) while still retaining 99.76% of the data, thus showing clear specificity towards discarding unreliable results. For the data at hand, and with mentioned filter settings (HWE, minimal read count of 10), MAGE yielded (almost) complete genotyping data (genotypes for at least 90% of the samples) for 143,069 SNPs, of which 134,971 (94%) not covered by the SNP array, confirming the merit of RNAseq-driven genotyping for subsequent ASE analyses.

**Figure 2:**
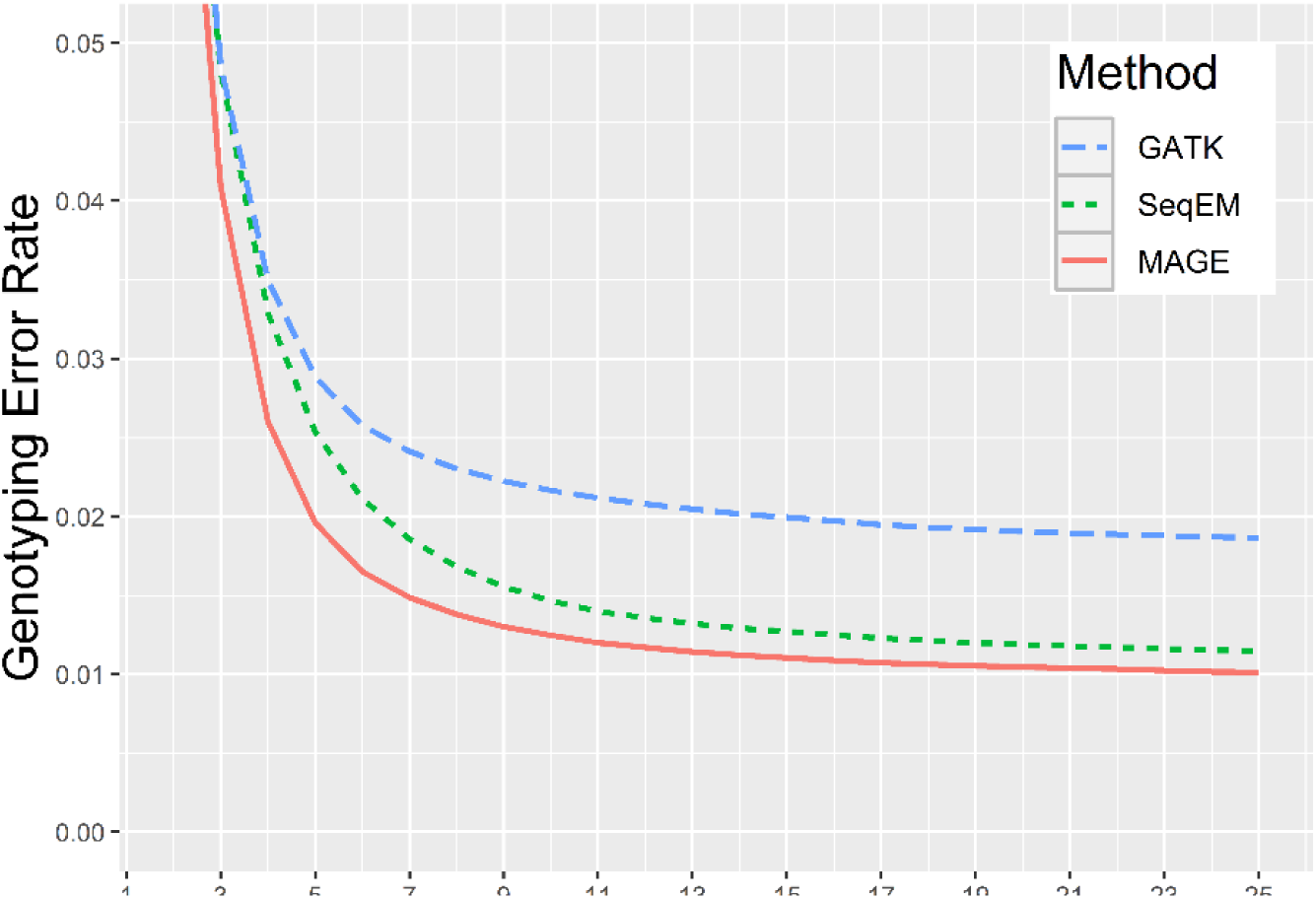
RNAseq-driven genotyping comparison – The genotyping error rate of GATK, SeqEM and MAGE for healthy kidney tissue (TCGA) RNA-seq data as a function of sequencing depth (per-SNP per-sample read count filter).

It should be noted that, in theory, also posterior genotype probabilities could be used for filtering. However, as the heterozygous fraction of samples is harder to model than the homozygous ones, posterior probability based filtering will particularly lead to removal of heterozygous samples or – when performed at the locus level – loci with larger fractions of heterozygous samples. While suboptimal for normal genotyping, this filtering strategy will be downright harmful for (d)AD analyses as one would be selectively removing the most informative data.

#### 2.2.B. (Differential) AD-detection in TCGA KIRC tumour and control data

Applying MAGE’s (differential) AD analysis (Section 2.1.B) on TCGA KIRC (tumour stage 1) and control data reveals a massive increase of AD in KIRC: of the 57,694 SNPs suited for AD-analysis (see methods), 8181 exhibited significant dAD at the 5% FDR level, with 7498 having increased AD in tumours. Summarizing these results to the gene-level a-posteriori (Section 4.2.F), 3444 of 11,496 genes considered feature significant dAD, with 3125 (91%) having increased AD in tumours. Given this large number of significant results, coupled with the fact that effect-size comparisons between individual loci/genes is complicated due to the interfering impact of tumour purity (see Discussion), we focus on chromosomal regions by means of rolling medians^19^. Top panels of Figure 3 show (d)AD and differential expression (DE) results for four chromosomes of interest (see Supplementary Section X.1 for all chromosomes), whereas bottom panels feature additional TCGA data (CNA, average gain and loss; DNA promoter hypermethylation, Section 4.2.E) to clarify the observed (d)AD results.

**Figure 3:**
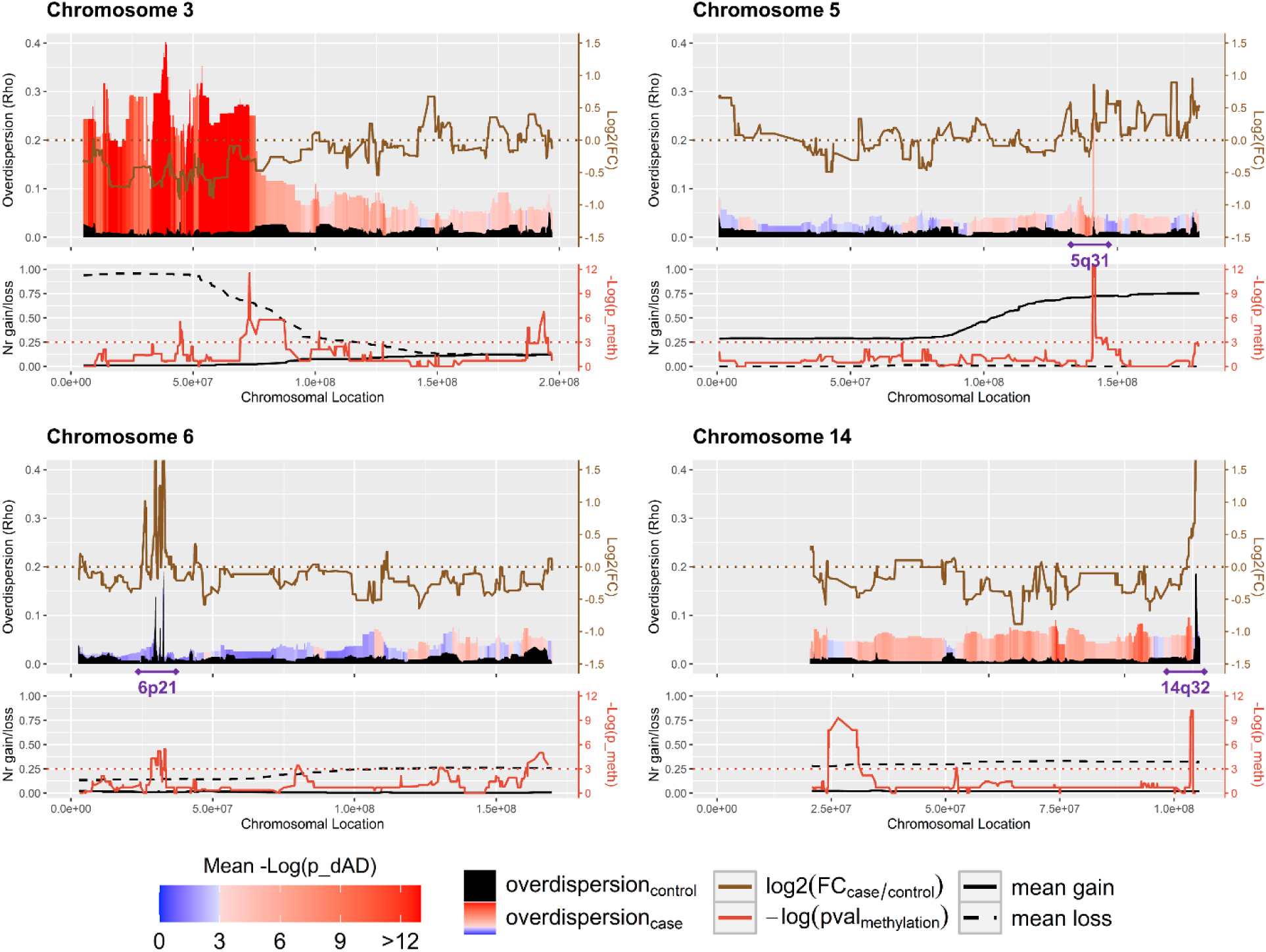
chromosome-wide (d)AD-results (area plots in top panels; case-ρ-values are color-coded according to statistical significance of dAD), differential expression log_2_[Fold Change] of expression (solid brown line in top panels; horizontal dotted brown line marks log_2_[Fold Change] = 0), case-hypermethylation fisher exact test p-values (solid red line in bottom panels, dotted red line represents 5% FDR) and average number of gain- and loss-events (solid and dashed black lines in bottom panels, respectively), visualized as per-gene results rolling averages/medians (window size = 15 genes; see Methods for details). Some regions of interest are marked in purple.

Even in healthy kidney, several regions characterized by a large AD are identified (overdispersion control, black). These are known to feature RME, e.g. HLA^20^ (6p21) and immunoglobulin variable chain^21^ (IGHV, 14q32) regions (Figure 3). These regions typically don’t exhibit dAD, yet feature overexpression in cancer, which can be explained by the rolle of (neo)angiogenesis in cancer, especially since the corresponding RME genes are expressed in leukocytes^20,21^.

Supported by TCGA CNA data but also RNA-seq DE-results, KIRC dAD results predominantly coincide with chromosomal aberrations in cancer, including the common 3p deletion, 5q gain, 14q loss (Figure 3), 7 gain, and 16 gain events^22^. However, dAD in KIRC goes beyond CNA, as demonstrated for the 5q31 region, which features at most minor AD in controls, but clearly significant dAD in cancer. The 5q31 contains several protocadherin (PCDH) gene clusters, which were demonstrated to feature long-range epigenetic silencing in cancer.^23^ The latter is also the case in KIRC (DNA methylation, Figure 3) and MAGE clearly indicates that silencing acts on individual alleles, contrasting other genomic regions featuring hypermethylation.

The same 5q31 region is part of a larger region often gained in KIRC (CNA gain results Figure 3), anticipated to have the opposite effect of methylation on expression. When evaluated across 21 PCDH cluster genes on 5q31, the methylation-expression correlation was clear (nine significant Spearman rank correlations, 5% FDR, mean correlation -0.19), but the CNA-expression correlation less so (only two significant correlations, mean correlation 0.13). Interestingly, CNA gains and DNA hypermethylation occurred independently (zero significant correlations, mean correlation 0.04) and in samples with simultaneous CNA gain and hypermethylation no indication of preferred silencing of the gained or non-gained allele was found (Supplementary Section X.2). The Discussion explores the implications of this apparent lack of coordination.

Note that for the results depicted in Figure 3, MAGE performed a joint control-case model fit with separate *ρ*_*het*_ for the purpose of dAD detection via LRT (Section 4.1.C). However, MAGE can also estimate *ρ*_*het*_ on control (or case) data separately, e.g. for RME studies in controls, yielding very similar results (Supplementary Section X.3).

#### 2.2.C. Detecting (loss of) imprinting in TCGA kidney data

Running MAGE’s monoallelic expression analysis (Section 2.1.C) on kidney control data and summarizing SNP results to the gene level (Section 4.2.F) yields 20 candidate imprinted genes (Table 1, 5% FDR level, estimated degree of (median) imprinting ≥ 0.85, see Section 4.2.C for a description of filtering settings). These include known imprinted genes such as *H19, MEG3* and *MEST*, but also novel candidates, some of which are likely false positive due to imprinting-independent monoallelic expression (e.g. *MIR6891*, located in the *HLA* gene cluster^12^). Subsequent evaluation of significant loss of imprinting and differential expression of these candidate imprinted genes (excluding *MIR6891*) identified eight genes showing increased biallelic expression in cases, of which four were associated with expression upregulation (log_2_FC cut-off of 0.5), i.e. canonical loss of imprinting: *HM13, H19, MEG3* and *U2AF1L4* (Table 1).

**Table 1:** MAGE’s candidate imprinting results, i.e. genes detected as putatively imprinted (Section 2.1.C, 5% FDR) with subsequent p-values assessing statistical significance of loss-of-imprinting (p (DI)) and DE (p (DE)), as well as log_2_FC and mean estimated imprinting across SNPs (î). N denotes the number of SNPs on which these gene-level results were based (Section 4.2.F). Significant p-values are marked in bold (5% FDR), gene symbols in bold are compatible with canonical LOI (both p-values < 0.05 and log_2_FC > 0.5; excluding MIR6891 from the HLA gene cluster). For genes marked with an asterisk, no gene-level expression data was available in the htseq-file provided by TCGA, and differential expression was evaluated by combining t-tests on their raw SNP-level counts (Section 4.2.F).

| Gene | N | p (DI) | p (DE) | Log2FC | $\hat{i}$ | Gene | N | p (DI) | p (DE) | Log2FC | $\hat{i}$ |
| --- | --- | --- | --- | --- | --- | --- | --- | --- | --- | --- | --- |
| *ZDBF2* | 6 | 8.52E-02 | **3.43E-15** | -0.96 | 0.98 | *PWAR6\** | 4 | 7.86E-01 | **1.91E-29** | -1.14 | 0.99 |
| *MIR6891\** | 1 | **4.70E-04** | **2.11E-19** | **2.33** | 0.94 | *SNHG14* | 15 | 3.02E-01 | **1.41E-09** | -0.56 | 0.97 |
| *COPG2IT1\** | 5 | 5.09E-01 | **2.62E-24** | -1.61 | 0.94 | *IPW\** | 2 | 2.62E-01 | **7.93E-18** | -0.72 | 0.99 |
| *MEST* | 2 | **6.02E-06** | **6.72E-11** | -1.15 | 0.9 | *ZNF597* | 5 | 3.32E-01 | **3.19E-02** | 0.16 | 0.95 |
| *GTF2IRD2* | 1 | 9.87E-01 | **9.16E-03** | 0.26 | 0.9 | *LINC01928\** | 1 | **2.41E-03** | **4.59E-04** | -1.24 | 0.89 |
| *LOC105375268\** | 1 | **3.33E-02** | **5.77E-18** | -1.71 | 0.86 | *PEG3* | 6 | 1.96E-01 | **7.03E-66** | -2.35 | 0.98 |
| *RP11-478H13.4\** | 2 | 5.28E-01 | **3.69E-05** | -1.36 | 0.9 | ***U2AF1L4*** | 1 | **8.09E-03** | **6.12E-19** | **1.14** | 0.85 |
| ***H19*** | 2 | **7.76E-06** | **1.20E-06** | **1.81** | 0.93 | *ZNF331* | 3 | **8.27E-18** | **1.36E-33** | -1.2 | 0.97 |
| ***MEG3*** | 7 | **2.44E-04** | **2.60E-02** | **0.56** | 0.95 | ***HM13*** | 3 | **1.24E-02** | **1.02E-16** | **0.56** | 0.97 |

## 3. Discussion

Besides zealous use in differential gene expression studies, RNAseq allows to discern between individual alleles, a feature that remains poorly exploited. Previous work allowed for RNAseq-driven genotyping based on EM^18^, yet did not take into account ASE. Similarly, beta-binomial based modelling of allele counts has already been proposed, yet overdispersion is either treated as a global nuisance hyperparameter to identify relevant *cis*-eQTLs^24^ rather than a locus-specific source of information, or methods focus on individual-level data of easily discernible heterozygotes instead of population-level genetics^25^. Though other RNAseq-only methods exist, they focus specifically on either *cis*-eQTLs^26^ or CNA^27^, not on AD phenomena such as imprinting, RME or dAD. MBASED^13^ is an exception, yet is solely tailored towards disease-associated EM and relies on individual (or paired) samples, forgoing the robustness of population-level analyses. Therefore, the Modeller of Allelic Gene Expression MAGE is the first exclusively RNAseq-based comprehensive AD-screening methodology. To this end, MAGE fits a beta-binomial mixture model to population-level RNAseq data, simultaneously genotyping the component individuals and capturing different phenomena such as AB (*cis*-eQTLs, alignment bias) and AD (methylation, CNA) in distributional parameters. These fits can subsequently be compared to null hypothesis models or fits on entirely different populations for differential AD studies via likelihood ratio tests. Imprinting and other AD phenomena causing complete monoallelic expression in controls are not easily captured by the fitted model. These are thus detected by a separate likelihood-based procedure while its dysregulation in cases is tested for by logistic regression.

Removing the reliance on DNA-based genotyping assays greatly improves cost- and data-efficiency. Indeed, as ASE-studies require large numbers of (heterozygous) individuals, population sizes should be sufficiently large to study loci featuring low minor allele (and hence heterozygote) frequencies, making DNA-based genotyping a major additional cost and a potential source of artefacts in case of genotyping errors. Moreover, limiting sequencing coverage or restricting the SNPs under analysis by exome-seq or the use of SNP arrays leads to far less loci under study, even when quality RNAseq data for those loci are available. For example, in the study at hand, 94% of the SNPs featuring high-quality RNAseq data were not available on the used SNP array. Consequently, the proposed RNAseq-only strategy allows to study AD in the large and ever increasing number of already available population-scale RNAseq datasets.

However, MAGE still has some limitations. First, RME detection via the imprinting-pipeline is only possible in case of imperfect mixture of cells expressing opposite alleles; only single-cell analysis can unequivocally identify all genes featuring RME. Moreover, MAGE avoids using large and costly genotyping assays but, being a population-level modeller, still requires large datasets. Smaller populations make genotyping and ASE-studies less reliable, but also exclude SNPs featuring low minor allele frequencies from further analyses. The assumed mixture model (Equation 1) has implicit limitations as well, a first one being the restriction to two alleles – or pairwise comparisons – per locus. While not problematic for human SNP-level data for which curated databases exist, it can be a drawback for studies pursuing completeness or being conducted on non-human organisms. Beta-multinomial models may offer a solution, yet suffer from more complicated model fits and the requirement of even more data. A second model limitation is that MAGE’s dAD analysis does not take into account sample impurity, which is particularly important in cancer, with KIRC as a clear example^22^. Given the impact on the dAD-effect size (change in overdispersion), especially for genes featuring high expression in the diluting non-tumour tissue, dAD-effect-size-based ranking is less appropriate to single out the most important genes. Therefore, here, we particularly focused on chromosomal regions featuring dAD rather than individual genes. Next to model-based adjustment for tumour impurity, which is statistically and computationally not straightforward, this problem may be solved by expanding our methodology to single-cell RNAseq data. Nevertheless, for non-cancerous phenotypes, it is anticipated that there will be substantially less aberrations, making gene prioritization far less of a problem.

Model limitations aside, there are additional data sources and analyses that, when incorporated in MAGE, would result in even more comprehensive ASE-studies. While MAGE currently focuses on SNP-level results, with post-analysis summary to the gene level, direct modelling at the transcript^28^ or gene-level and incorporation of additional small genetic variation such as indels could further improve power and interpretability, similar to Xie *et al*.^29^. Such a strategy could start from RNAseq-based phasing, e.g. using phASER^30^, yet it should be noted that state-of-the-art phASER-based allelic expression analyses typically still rely on DNA-based genotyping data^31^. Similarly, MAGE inferred genotypes can in principle be used for subsequent *cis*-eQTL studies, as the detection of AB during genotyping indicates presence of a *cis*-eQTL, yet alignment bias provides an alternative explanation. Though the latter could be minimized during preprocessing, as performed by the GATK RNAseq variant calling pipeline^32^ or the WASP^15^ remapping method, some amount of bias will most likely remain^33^. Moreover, these methods rely on specific assumptions and filter out different proportions of sequencing reads, implying that additional validation is required to pinpoint the optimal (MAGE-based) *cis*-eQTL detection strategy. Finally, it should be noted that additional experiments may be required to elucidate the (clinical) implications of obtained results. For example, cancer-related clonal expansion could lead to the mislabelling of random monoallelically expressed loci as dAD, and for some loci it remains unclear whether their monoallelic expression relates to imprinting or not.

The case study on renal cancer data proved MAGE’s superior RNAseq-driven genotyping over previously established methods. Subsequent dAD-analyses indicated large-scale genome-wide allelic dysregulation in KIRC, though very localized in specific genomic regions (which may hold valuable biomarkers) while other regions feature little to no dAD (Figure 3, Supplementary Results X.1). Of special interest are the protocadherin gene clusters at 5q31, for which MAGE demonstrates that the long-range epigenetic silencing observed in KIRC (yet previously also in other tumour types^23^) is in fact allele-specific. However, copy-number gains of this region were frequent as well, and associated with on average higher expression. Presence of both types of counteracting aberrations featured little correlation, neither negative nor positive, obscuring how exactly this region may be important in KIRC. One explanation is that particularly genes centromeric or close to the protocadherin clusters yield advantage to the 5q gain, with the protocadherin cluster getting caught up in the fray, explaining the lack of correlation. Finally, MAGE returned evidence of canonical LOI (i.e. associated with expression upregulation) for four genes in KIRC (*H19, MEG3, HM13* and *U2AF1L4*). In the past, *H19* and *MEG3* have been attributed both oncogenic and tumour suppressor properties^34–37^, making additional studies on the impact essential. Meanwhile, current and previous LOI results in breast cancer^16^, as well as functional studies^38,39^, support *HM13* as a candidate oncogene. *U2AF1L4* has been far less studied but is also of interest, being a homologue of a gene imprinted in mouse^40^ and featuring prognostic value in renal clear cell carcinoma according to preliminary experimental research^41,42^.

In conclusion, this study introduced the MAGE toolbox for RNAseq-driven genotyping and subsequent detection and statistical inference of AD-phenomena. Its merit was illustrated on KIRC data, outperforming previous RNAseq-driven genotypers and enabling genome-wide mapping of several ASE-phenomena. As an open source R software package, it is applicable in comprehensive studies on both existing and future RNAseq datasets to detect complex transcriptional responses beyond mere quantitative gene expression.

## 4. Methods

### 4.1. Mathematical implementation

MAGE’s mathematical procedures are available as an R programming language^43^ software package, though some subroutines use C and C++ to increase computational speed. This section provides a more in-depth description, while the package itself contains a vignette going over the entire pipeline as well (https://biobix.github.io/MAGE/articles/MAGE_tutorial.html).

#### 4.1.A. Hyperparameter estimation

Many of MAGE’s analyses revolve around fitting Equation 1 (Section 2.1.A) to allele counts using EM:

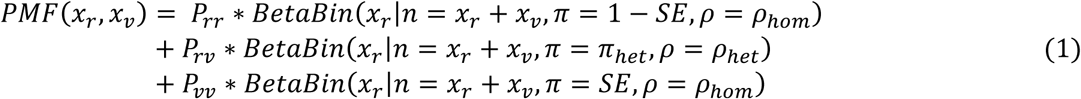

Before doing so, two hyperparameters need to be determined: the sequencing error rate (SE) and inbreeding coefficient (F; important in later HWE-filter calculations). While these can be set based on the sequencing methodology’s expected error rate (plus misalignment errors) and population (breeding scheme) knowledge, respectively, they can be roughly estimated from the provided RNAseq data as well, which may be mathematically more appropriate or provide a double-check. MAGE does so by fitting a simplified Equation 1 to all loci in the RNAseq data, using regular binomial mixture components without AB (π_*het*_ = 0.5) to get per-locus estimates of SE and F, the latter via EM-obtained preliminary genotyping results, as:

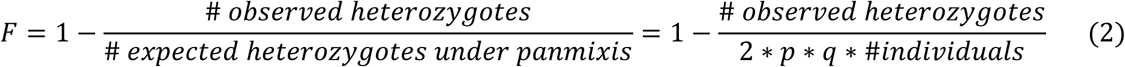

with p and q the observed reference- and variant-allele frequencies, respectively. As input loci should be plentiful, only high-fidelity loci are retained to calculate final median hyperparameter estimates (at least 10 samples with a median coverage of at least 4), besides requiring a sufficient minor allele frequency for reliable inbreeding estimation (> 0.15) and disregarding loci returning unreasonably high SE-estimates (< 0.035). Note that setting SE extremely low, though not necessarily unrealistic, can be risky, as this may lead to only pure reference- or variant-allele containing samples being recognized as homozygotes in subsequent EM procedures. Using a minimal lenient SE, such as 0.002-0.003, provides robustness to further analyses.

#### 4.1.B. Genotyping & AB detection

Even though Section 2 introduced the beta-binomial overdispersion parameter as *ρ*, ranging from zero to one, MAGE’s code uses an alternative parameterization *θ*, ranging from zero to infinity. These are merely parameter transformations of one another (Equation 3):

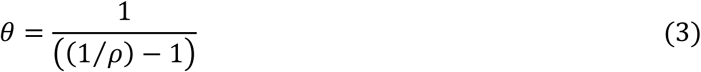

*ρ* is more practical for reporting results due to its zero-to-one range (e.g. visualizations like Figure 3) while *θ* often leads to simpler mathematical expressions of the beta-binomial distribution’s properties. To reflect MAGE’s actual code, *θ* is used instead of *ρ* throughout the remainder of this subsection.

Fitting Equation 1 using EM provides MAGE’s genotyping and AB detection functionalities (Section 2.1.A). Here we provide additional details regarding the specific EM implementation at hand, i.e. for fitting a beta-binomial mixture model to homozygous and heterozygous allele counts:

1. While Equation 1’s mixture components (*P*_*rr*_, *P*_*υυ*_, *P*_*rυ*_) receive balanced starting values of (1/3), a moment estimator is used for beta-binomial-specific starting parameters. This is non-trivial for observations with varying *n*, but based on *Kleinman*^44^ we derived:

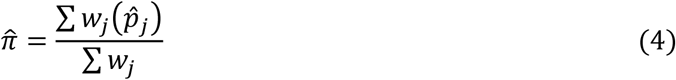

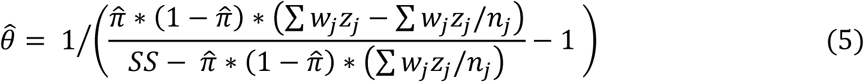 With 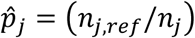 the fraction of reference-reads in sample *j*, 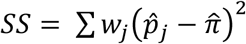 a weighted sum-of-squares, *z*_*j*_ = (1 − *w*_*j*_/∑ *w*_*j*_), and *w*_*j*_ a per-sample weight between 1 and *n*_*j*_ for which *Kleinman*^44^ suggests an iterative procedure. However, given our aim to find a rough initial estimate, all *w*_*j*_ are simply set to 1, which corresponds to the ideal weights for a *θ* = 0 scenario.
2. Though there is no shortage of beta-binomial PMF implementations in R, these proved to be either slow or return nonsensical results when pushed to extreme parameter values (or default to binomial densities in the case of *VGAM*^45^). While extreme parameters are not necessarily realistic, numerical optimization algorithms can encounter them while exploring the parameter space, leading to MAGE’s own PMF implementation to ensure numerical stability. This is either based on a representation as long products^46^ (implemented in C):

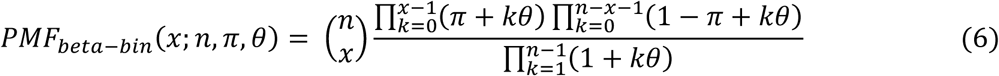

or using beta-functions (written as gamma-functions for C++ implementations):

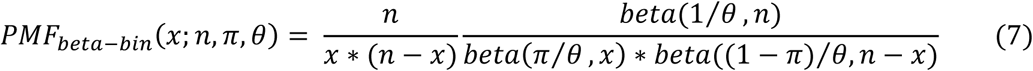 The former is faster for lower-count samples (up to about 50 reads), the latter for higher-count samples, and MAGE switches between the two accordingly. To avoid numerical precision problems, all PMF calculations happen log-transformed; despite this, extreme parameter values can still yield extreme *log-beta* values and subsequent catastrophic cancellation in the final result of Equation 7, in which case MAGE switches to a C++ implementation so as to use the boost multiprecision library^47^ to avoid the issue.
3. MAGE’s EM includes maximum likelihood estimation of Equation 1’s parameters, which occurs via numerical optimization relying on log-likelihood (and gradient) functions. This is a bounded problem (π ∈ [0,1], *θ* ∈ [0, ∞[), yet bounded optimization algorithms, such as *optim*’s L-BFGS-B (base R), proved numerically unreliable. Therefore, optimization happened unconstrained using *optim*’s BFGS algorithm on transformed parameter values, i.e. a logit- and log-transform on π and *θ*, respectively.
4. Beta-binomial PMFs are bimodal when *θ* > max(π, 1 − π). Such bimodality is undesirable for allele count data already modelled as a mixture, so MAGE redoes parameter estimation using *alabama*’s Augmented Lagrangian algorithm^48^ if this occurs. This allows for non-linear constraints in parameters (*θ* > max(π, 1 − π)), but is considerably slower than *optim*’s BFGS, hence why it is not MAGE’s default option.
5. MAGE’s EM is robust, incorporating outlier detection based on case-deletion MLEs first proposed by Cook^49^. For each locus, let 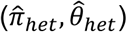 be the final full-data estimates of both heterozygous beta-binomial parameters and 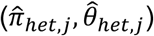 re-fitted parameter estimates with the *j*^*th*^ sample deleted. Then, 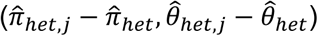 are two general measures of sample *j’s* influence on parameter estimation. Once calculated for all samples, those for which either of these measures deviates more than 5 (default value) sample standard deviations from the sample mean are considered high-impact outliers, and the entire analysis is re-done on non-outlying samples. In practice, outlier removal is not that impactful for genotyping performance while having a considerable computational cost, so one may decide against it. It is very important when performing statistical inference on the *θ*-parameter though (dAD-detection) as overdispersion-estimation is very sensitive to high-leverage outliers. Even though outlying samples play no part in the model fit, they are still genotyped using said model and remain included in the final results (marked as outlier). Besides completeness, this is important for unbiased assessment of HWE on genotyping results (see point 7) as outlier detection, being performed on the heterozygous parameters, mainly removes (high-leverage) heterozygous samples.
6. Significant AB is detected by re-doing the entire fit on the (outlier-free) dataset with π_*het*_ set to 0.5, then performing an LRT (test statistic 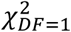).
7. HWE-conformity is assessed after genotyping, by comparing each locus’ fitted genotype counts 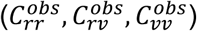 with those expected under HWE via base R’s *chisq*.*test* (using the previously estimated hyperparameter *F*):

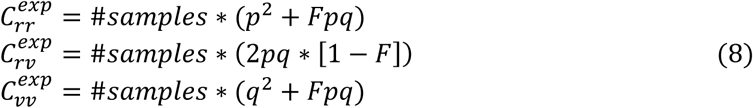

With:

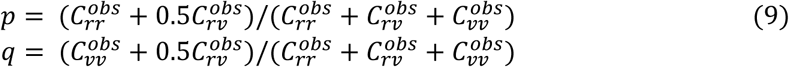 In some cases, e.g. specific breeding designs, *F* can be negative (outbreeding). For one specific locus and assuming *q* is the minor allele frequency, the theoretical per-locus minimum of *F* equals −*q*/(1 − *q*), as lower values of *F* yield negative expected genotype counts. Since *F* is estimated as a locus-wide median, it can indeed be lower than some locus’ theoretical minimum, in which case that locus’ HWE-conformity test is performed using *F* = −*q*/(1 − *q*) instead. A locus is considered HWE-conform if its *chisq*.*test* p-value is greater than 0.001, a (somewhat arbitrary) cut-off that is nevertheless standard^50–52^.

#### 4.1.C. dAD analyses

Before assessing dAD, Equation 1 is fit separately to both control- and case-datasets for the sole purpose of outlier-detection (Section 4.1.B); it’s inappropriate to perform outlier-detection during the two dAD-detecting joint fits as (1) these fits need to be performed on the same input datasets for valid LRTs, and (2) outlier detection based on case-deletion MLEs 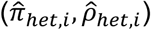 implicitly assumes a single underlying distribution, thus with as single true *ρ*_*het*_, while identifying differences in *ρ*_*het*_ between cases and controls is the main goal here.

After outlier detection, MAGE fits Equation 1 to the joint case and control dataset and compares it to a fit sharing all parameters between cases and controls except *ρ*_*het*_:

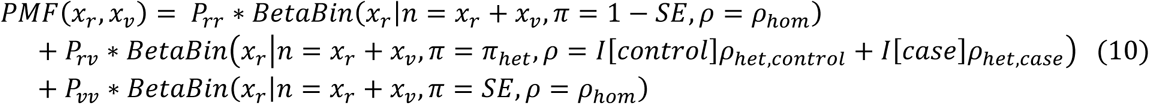

With *I*[*control*] and *I*[*case*] indicator variables for each sample’s population. An LRT comparing fits of Equations 1 and 10 (test statistic ∼ 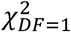) detects significant dAD. Remarks from Section 4.1.B typically apply to dAD-analysis as well.

#### 4.1.D. Imprinting analyses

(Differential) imprinting analysis relies on basic concepts introduced earlier^16^, yet with several improvements summarized here.

To speed up imprinting detection (on control datasets), knowing that imprinting is commonly rather extreme (i.e. heterozygotes will mainly express one of their alleles, being apparently homozygotic in RNAseq data), loci are first passed through a symmetry filter testing whether the observed number of samples with reference allele ratios greater and lesser than 0.5 are proportionate to the observed reference- and variant allele frequencies, respectively (via R’s *chisq*.*test*). This holds true regardless of the inbreeding coefficient *F*; designating the observed reference- and variant allele frequencies as *p* and *q*, respectively and assuming a heterozygote’s alleles have an equal chance to be imprinted, the expected numbers are:

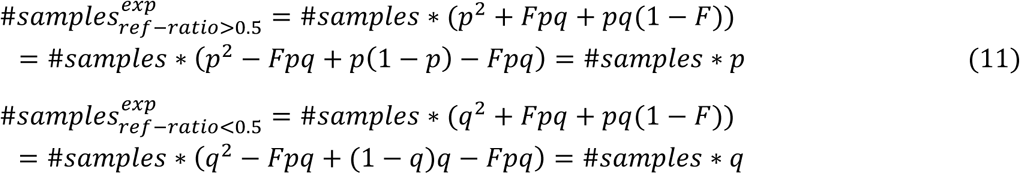

For loci passing the symmetry filter, data likelihoods are calculated according to the following PMF, with *i* varying from 0 to 1 in steps of 0.01:

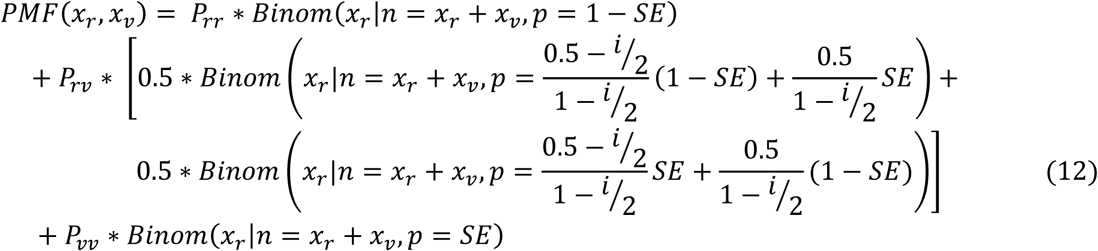

The most likely *i* is retained and tested for significant imprinting via LRT against the *i* = 0 fit (test statistic ∼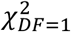). Subsequent LOI-detection, which has been improved from our previously published imprinting analysis^16^, then takes both inbreeding and sequencing depth into consideration via binomial logistic regression: every sample’s “degree of heterozygosity” can be defined as the read coverage of its least over its most expressed allele (1 for perfect heterozygotes, 0 for both perfect homozygotes). MAGE then models this per-sample ratio with respect to control-case status via binomial logistic regression (using R’s *glm* function with *family=binomial* from the base R *stats* package), with a significant and positive control-case regression coefficient indicating LOI. The latter is explicitly tested via LRT instead of *glm*’s default Wald test, since we noticed that the latter may numerically fail for the most obviously imprinted cases. Wald tests are indeed known to sometimes fail on extreme data^53^, making the more reliable LRT preferable.

#### 4.1.E. MAGE visualization

It should be remarked that the shape of beta-binomial distributions, even when rescaled to [0,1], still depends on the total read count *n*, which varies per sample. MAGE’s distribution visualizations (lineplots in Figure 1A) therefore use a locus’ median *n*. This implies that the figures are for illustrative purposes only, and lineplots don’t necessarily fit the underlying samples (histogram) that well even in case of a very good model fit, unless the latter are transformed to same-quantile observations using said median *n*.

### 4.2. Renal (cancer) case study

#### 4.2.A. Data acquisition

RNAseq BAM-files (GRCh38) were downloaded from TCGA^54^, of which 128 control kidney samples (72 KIRC, 32 KIRP, 24 KICH) and 534 KIRC tumour samples, 268 of those being stage 1 tumour samples used in our case study (both deduplicated; in case of duplicates the sample with the most recent timestamp was retained). To assess genotype performance, we downloaded Affymetrix Human SNP array 6.0 data (906,600 SNPs assayed, GRCh37) from the GDC legacy archive^55^ as well as the GenomeWideSNP_6 annotation file (release 35) accessible from Affymetrix^56^, to link Affymetrix ids to dbSNP-ids. Xenabrowser^57^ provided Infinium HumanMethylation 450k and CNA data to compare against dAD-results (Sections 2.2.B, 4.2.E). For genotyping, all kidney control samples were used whereas for (d)AD analyses, solely KIRC stage 1 samples and KIRC controls were considered.

#### 4.2.B. Data preprocessing

mpileup/bcftools (v0.1.9) calls variants from BAM files, after indexing if necessary^58^, retaining only those with a minimal raw read depth (default 10) in at least one sample and present in dbSNP^59^. By default, non-uniquely mapped reads are filtered out to reduce noise. Per-sample allelic counts for all possible variants (A/C/G/T) were written to count files together with dbSNP reference alleles (if available) and sample ID, to be used as input for subsequent analyses in R. Subsequently, MAGE first determines exactly one reference- and one variant-allele as the most frequent dbSNP reference alleles (percentage-wise across all loci; when no dbSNP reference is available all possible bases are treated as potential reference alleles). The final per-locus prior filter is the requirement of at least 10 samples covering one (or both) of this reference- and variant allele. The entire pre-processing pipeline starting from BAM files is available through Github, as well as R-scripts for all subsequent analyses using the MAGE package; a Conda environment file ensures reproducibility.

#### 4.2.C. ASE analyses and applied filters

All ASE analyses were performed as described in Sections 2.1, 2.2 and 4.1, with additional filter settings described below. To improve the dAD results’ qualtiy (Section 2.2.B), prior per-SNP filtering used these minimal criteria: HWE chi-square p-value > 0.001 (evaluated on control data), median coverage in tumours and controls ≥ 4, at least 12 estimated heterozygotes in control data. For imprinting detection (Section 2.2.C), besides similar generic filters (minimal median coverage > 4, at least 30 samples, passing the symmetry filter from Section 4.1.D), there’s a filter on each locus’ minor allele frequency, so as to retain loci on which imprinting can be reliably detected (a minimal minor allele frequency ensures a minimal number of expected heterozygotes). This happens robustly by requiring both an estimated minor allele frequency of at least 0.1 based on aggregated raw allele counts, and of at least 0.15 based on the simplified genotyping from the hyperparameter estimation procedure (Section 4.1.A).

Posterior filtering for imprinting-analyses happens after summarization to gene-level (Section 4.2.F). Besides statistical significance, we require a minimal effect size of *i* > 0.85, on top of a robust median imprinting of at least 0.85, both evaluated as gene-level weighted means. The latter is determined by sorting all samples of a locus according to the degree of heterozygosity (minor over major allele count), removing the number of expected – under HWE - homozygotes (i.e. the bottom #*samples* ∗ (*p*^2^ + 2*F*_*pq*_+*q*^2^) results), then calculating the degree of imprinting of the median minor over major allele ratio of the remaining samples as 2 ∗ (0.5 − (*minor allele count*/ *major allele count*)).

#### 4.2.D. GATK & SeqEM genotyping

Section 2.2.A evaluated MAGE’s RNAseq-driven genotyping performance against those of GATK and SeqEM. For the former, we followed GATK’s RNAseq Short Variant Discovery pipeline^17^ using default parameters, except for specifying *–output-mode EMIT_ALL_ACTIVE_SITES* in the *gatk HaplotypeCaller* command so as to make GATK call homozygous samples as it doesn’t do so by default. For SeqEM, we implemented the model described by the authors^18^ in MAGE ourselves. While SeqEM is also available as a Windows executable file, it was very time-consuming and performed worse than our own implementation, most probably due to numerical problems while fitting for several loci.

To allow for an unbiased comparison of genotyping performance, no filters depending on MAGE’s model fit are applied; every per-SNP per-sample genotype is retained if it passes the default Birdseed confidence score threshold^60^ (<= 0.5) in the Affymetrix data, the SNP is covered by 10 or more samples in the RNAseq data, and it is assigned a genotype by GATK’s pipeline using default filters.

#### 4.2.E. DE- and hypermethylation-detection

Section 2.2.B compared dAD results to DE-results, average CNA and tumour hypermethylation. For per-gene DE-results, we downloaded htseq gene count files from Xenabrowser^57^, which were processed in R using the *EdgeR*^1^ package. Xenabrowser provided CNA and methylation data as well, the latter of which was tested for significant hypermethylation in tumour-compared to control-samples via Fisher exact test on the number of highly methylated samples (defined as having a methylation percentage > 20% according to the most population-wide hypermethylated CpG in each gene’s promotor region; specific CpG determined separately for the control- and tumour-population).

DE results are also included in the imprinting analysis (Table 2) but, given the focus on individual genes instead of larger regions, the small number of significant LOI results, and the fact that some genes were missing expression data in the TCGA-provided htseq files, DE-results were obtained on a per-SNP basis using only the reference- and variant-allele count, via R’s *t*.*test* function on library size-corrected log count-per-million data when necessary (asterisks in Table 1). These were then combined into gene-level results (Section 4.2.F).

#### 4.2.F. SNP-to-gene summarization

Sections 2.2.B and 2.2.C include gene-level results, even though MAGE’s raw results are returned at SNP-level. This is achieved by combining results via either geometric means (p-values) or arithmetic means (overdispersion-parameters, estimated (robust median) imprinting, log_2_FC). Wilson^61^ discusses several ways of p-value combination – minimum, harmonic mean, geometric mean, arithmetic mean, maximum – which all make some trade-off and can be written as a general formula containing a variable *r* parameter. This formula provides even more nuance and is implemented in MAGE to customize p-value combination. We found the geometric mean to strike a good balance: it’s fair to expect most SNPs of a gene to provide evidence supporting dAD, (LO)I or DE. Minimum and harmonic mean seem overly liberal, as a single – e.g. false positive - significant p-value is typically sufficient to achieve overall significance. The arithmetic mean and maximum, on the other hand, are rather conservative as a single non-significant SNP – e.g. due to technical reasons – easily leads to overall non-significant results.

The per-SNP geometric and arithmetic means discussed above are weighted according to the square root of {median coverage times the number of expected heterozygous samples} per locus, the latter of which is either a result of Equation 10’s EM-fit (for dAD) or calculated according to HWE (for imprinted loci). For dAD, LOI and DE p-values, as well as log_2_FC, which are based on a comparison between a control- and case-population, the minimum of this weight across both populations is used; results calculated on one population (control- or case-overdispersion, degree of (robust median) imprinting in controls) use just the corresponding population to calculate a locus’ weight.

For SNP-annotation, we webscraped the ncbi database^62^ (https://www.ncbi.nlm.nih.gov/snp/) which returns a per-SNP list of candidate genes. We determined a “hierarchy” among these annotations (based on the likelihood to observe RNAseq-data for this annotation) and assigned each SNP to the gene highest on this hierarchy; in case of a tie, the SNP was not used for summarization. The hierarchy goes {Missense Variant | Synonymous Variant | Initiator Codon Variant | Terminator Codon Variant | Stop Gained | Stop Lost | Coding Sequence Variant} > {5 Prime UTR Variant | 3 Prime UTR Variant | Non Coding Transcript Variant} > {Intron Variant | Splice Acceptor Variant | Splice Donor Variant} > {2KB Upstream Variant | 500B Downstream Variant}. For SNPs missing a gene-annotation on ncbi, we tried finding a unique gene match using the R package *GenomicRanges*^63^, supported by *ensembldb*^64^, *GenomeInfoDb*^65^, *BSgenome*^66^, *SNPlocs*.*Hsapiens*.*dbSNP151*.*GRCh38*^68^ and *EnsDb*.*Hsapiens*.*v86*^67^.

## Supporting information

Supplementary Results

## 5. Acknowledgements

The results published here are based upon data generated by the TCGA Research Network: https://www.cancer.gov/tcga. This research was funded by the Research Foundation Flanders (FWO grant 1128021N) and BOF grants BOF.24Y.2019.0020.01 and BOF.DOC.2019.0036.02.

