## Supplementary Results for "MAGE enables population level RNAseq driven genotyping and (differential) allelic divergence detection in healthy kidney and carcinoma"

### X. Supplementary results

#### X.1. dAD-plots of all chromosomes

**Chromosome 1**

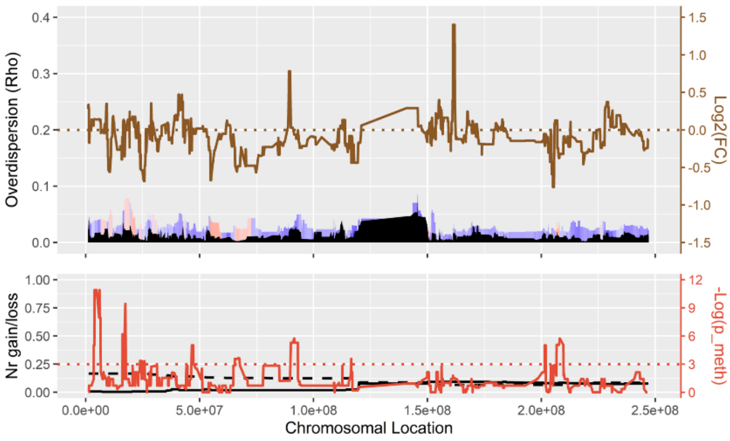


**Chromosome 2**

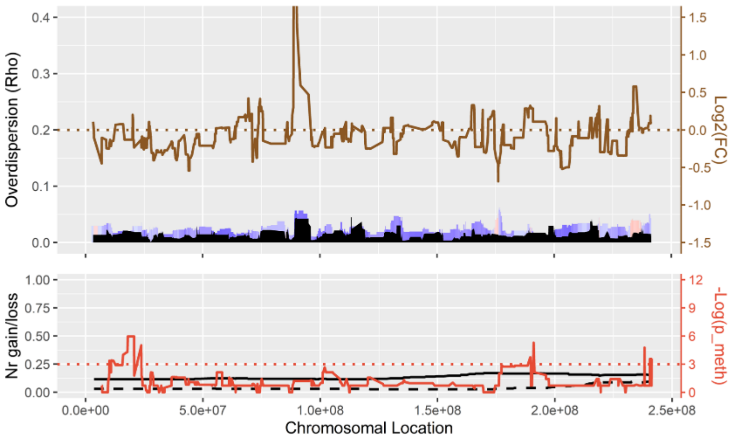


**Chromosome 3**

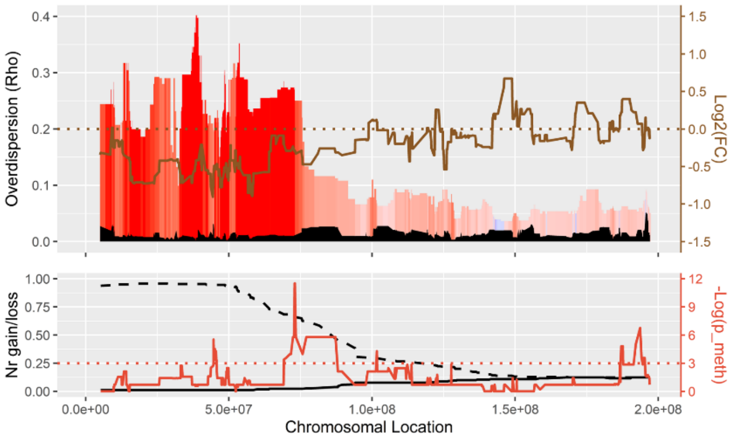


**Chromosome 4**

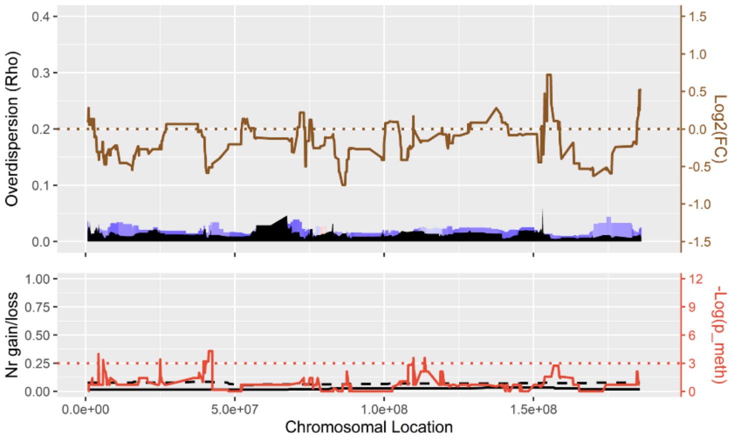


**Chromosome 5**

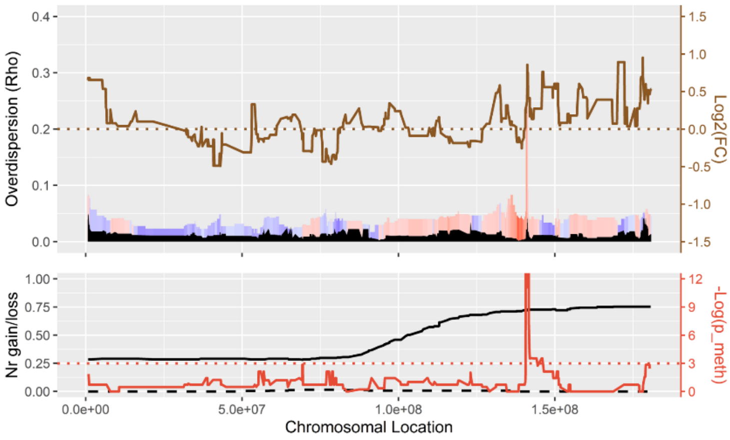


**Chromosome 6**

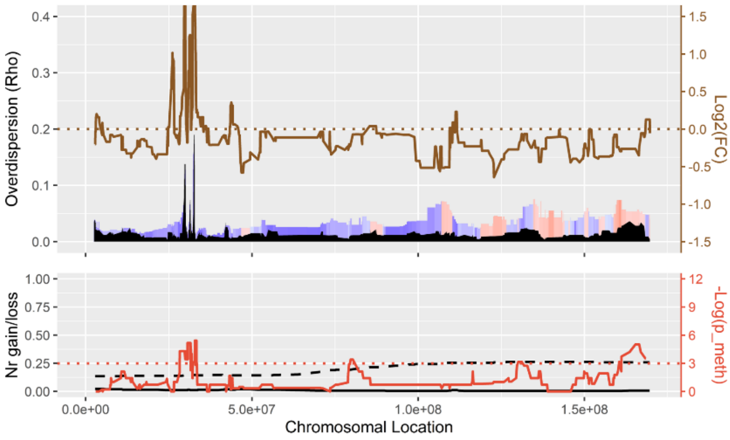


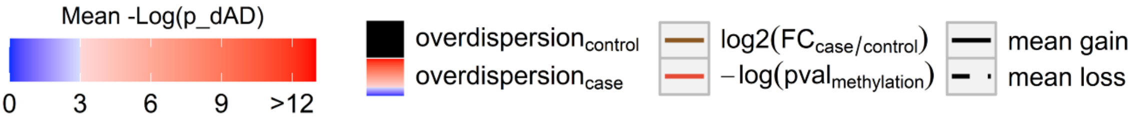

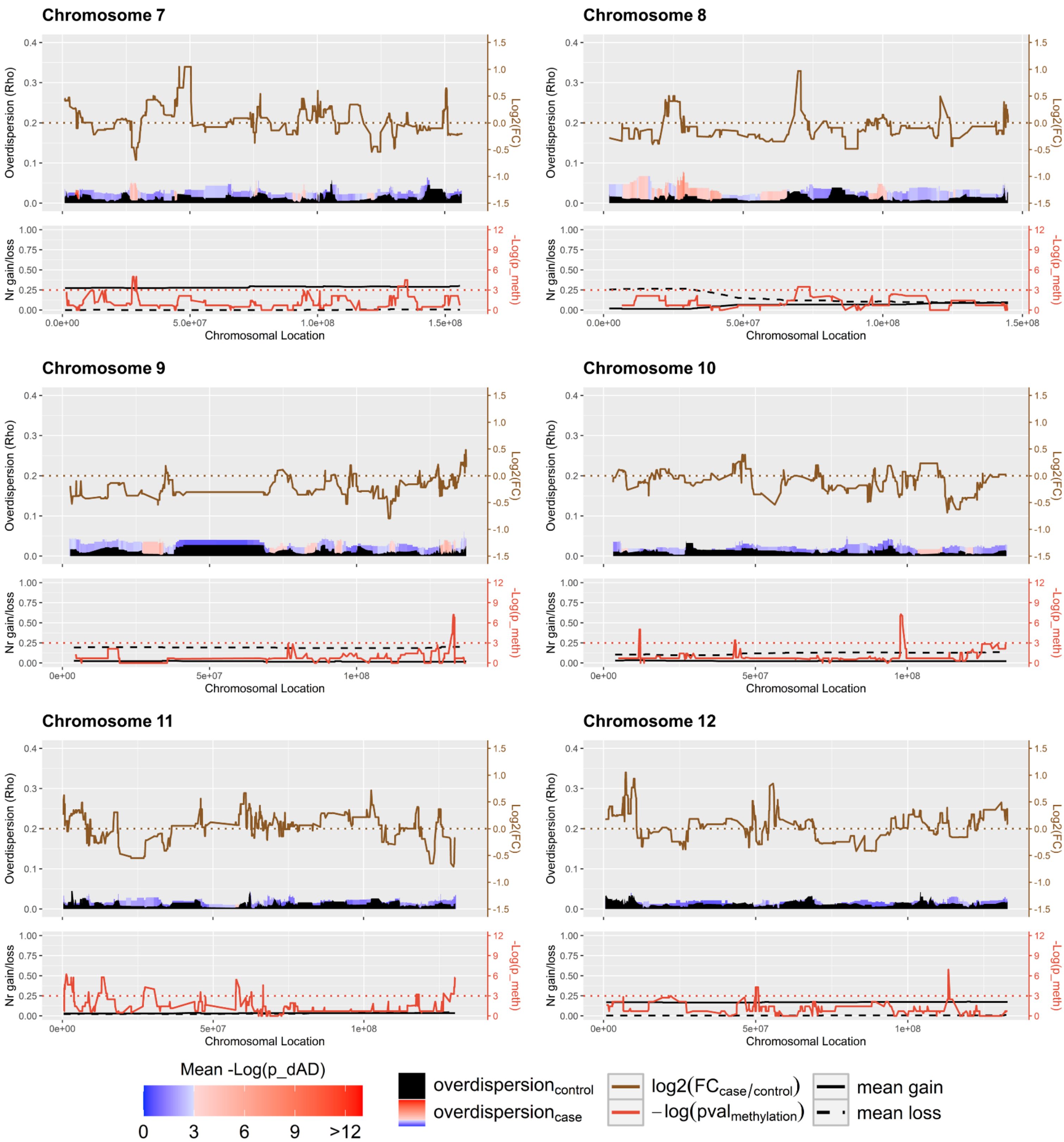

Chromosome 7
Chromosome 8
Chromosome 9
Chromosome 10
Chromosome 11
Chromosome 12
Overdispersion (Rho)
Log2(FC)
Nr gain/loss
-Log(p_meth)
Chromosomal Location
Mean -Log(p_dAD)
0
3
6
9
>12
overdispersion_control
overdispersion_case
log2(FC_case/control)
−log(pval_methylation)
mean gain
mean loss

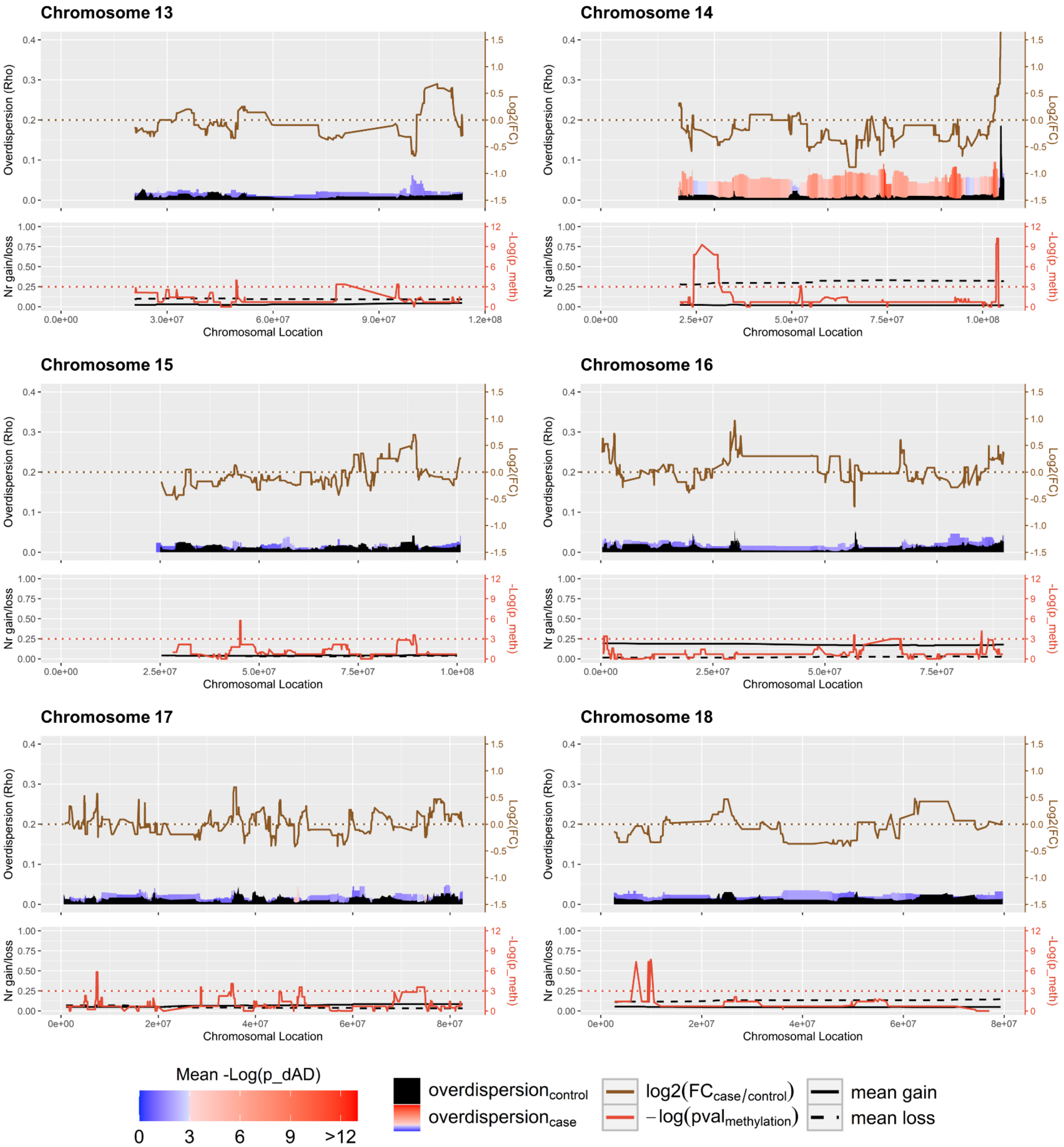
Chromosome 13
Chromosome 14
Chromosome 15
Chromosome 16
Chromosome 17
Chromosome 18
Overdispersion (Rho)
Log2(FC)
Nr gain/loss
-Log(p_meth)
Chromosomal Location
0.0e+00
3.0e+07
6.0e+07
9.0e+07
1.2e+08
2.5e+07
5.0e+07
7.5e+07
1.0e+08
0e+00
2e+07
4e+07
6e+07
8e+07
Mean -Log(p_dAD)
0
3
6
9
>12
overdispersion$_{control}$
overdispersion$_{case}$
log2(FC$_{case/control}$)
−log(pval$_{methylation}$)
mean gain
mean loss

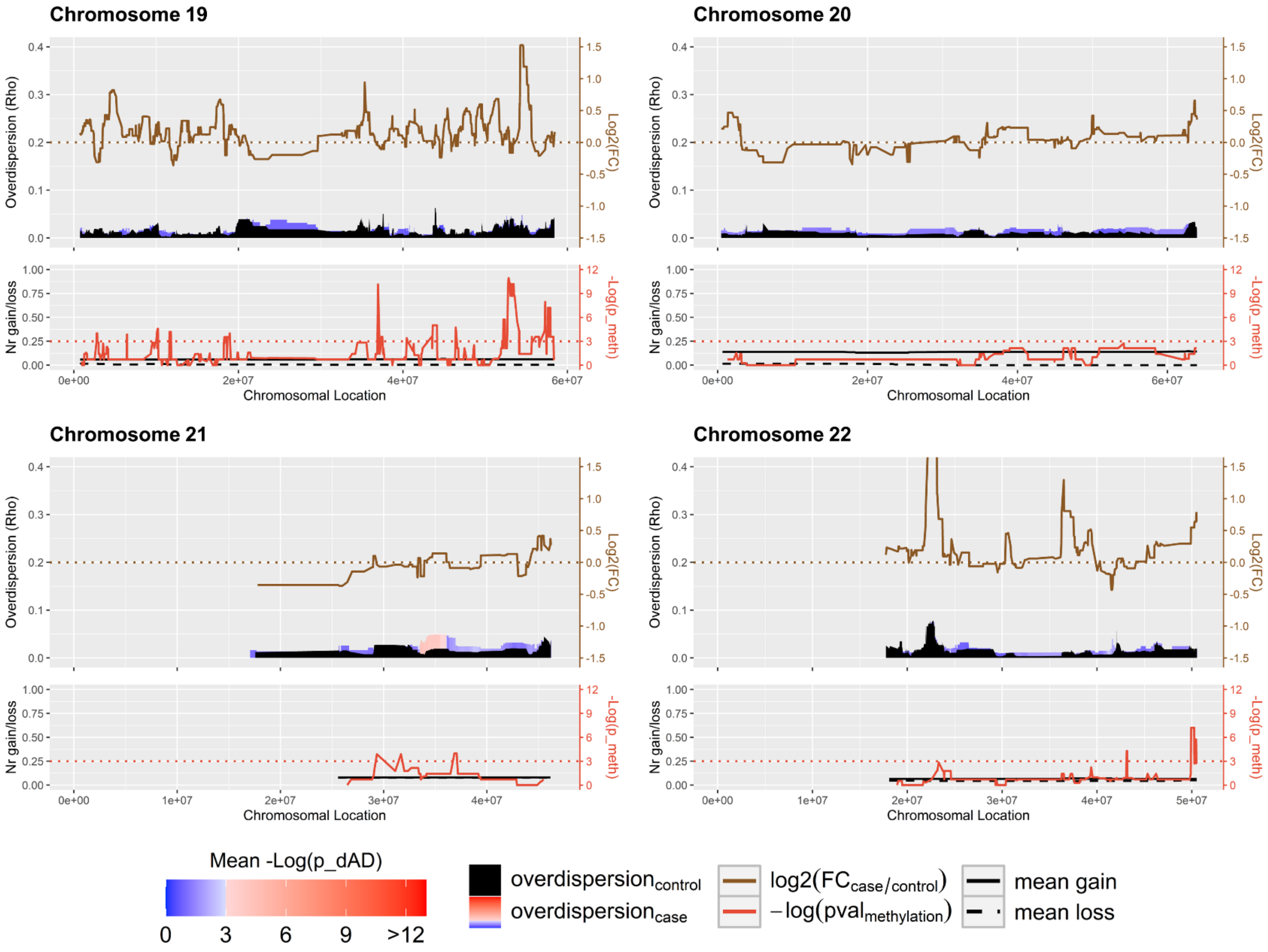

Chromosome 19
Chromosome 20
Chromosome 21
Chromosome 22
Overdispersion (Rho)
Log2(FC)
Nr gain/loss
-Log(p_meth)
Chromosomal Location
Mean -Log(p_dAD)
0
3
6
9
>12
overdispersion_control
overdispersion_case
log2(FC_case/control)
−log(pval_methylation)
mean gain
mean loss

#### X.2. Exploratory CNA/methylation plots of PCDH genes

This Section contains several exploratory reference allele fraction histograms of SNPs lying in genes in PCDH (protocadherin) clusters genes, each with a scatterplot panel underneath showing whether the samples (depicted in the histogram) exhibit either CNA (copy number gains, red), hypermethylation (>20% for the most differentially hypermethylated CpG in the promoter region of the gene containing the SNP; blue), both (purple) or neither (grey). Samples for which this information is incomplete are not labeled. These plots allow to explore whether e.g. in samples featuring both copy number gains and promoter hypermethylation, particularly the gained allele is silenced by hypermethylation to avoid excess expression of the PCDH cluster genes (leading to allele fractions closer to 0.5, antagonistic effect), whether a synergistic effect is present by particularly silencing the non-gained allele (leading to allele fractions closer to 0 to 1).

These plots were evaluated for 30 SNPs (four depicted here), but for none of them any indication was found of CNA and hypermethylation working in a concerted manner on relative allelic expression. Visually, the samples labeled purple (both CNA and hypermethylated) lie neither consistently more towards the extremes (reference allele fraction of 0 or 1; synergy) than the red and blue dots or vice-versa (antagonism), thus not conveying any clear coordination between these phenomena (see Sections 2.2.B, 3).

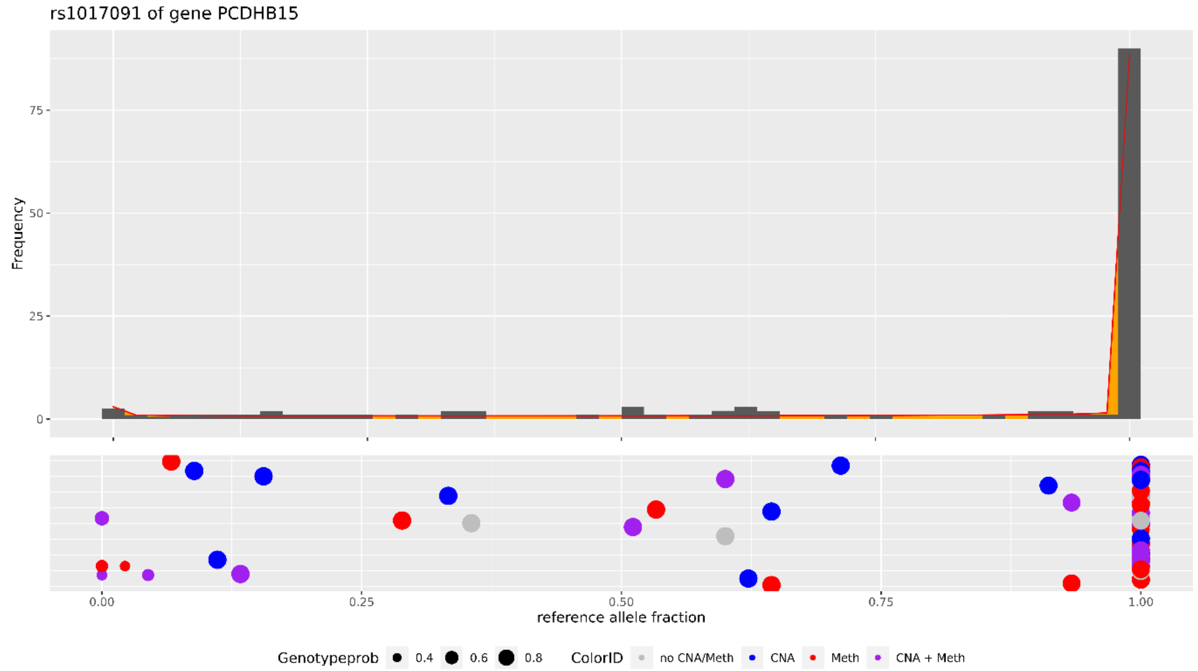

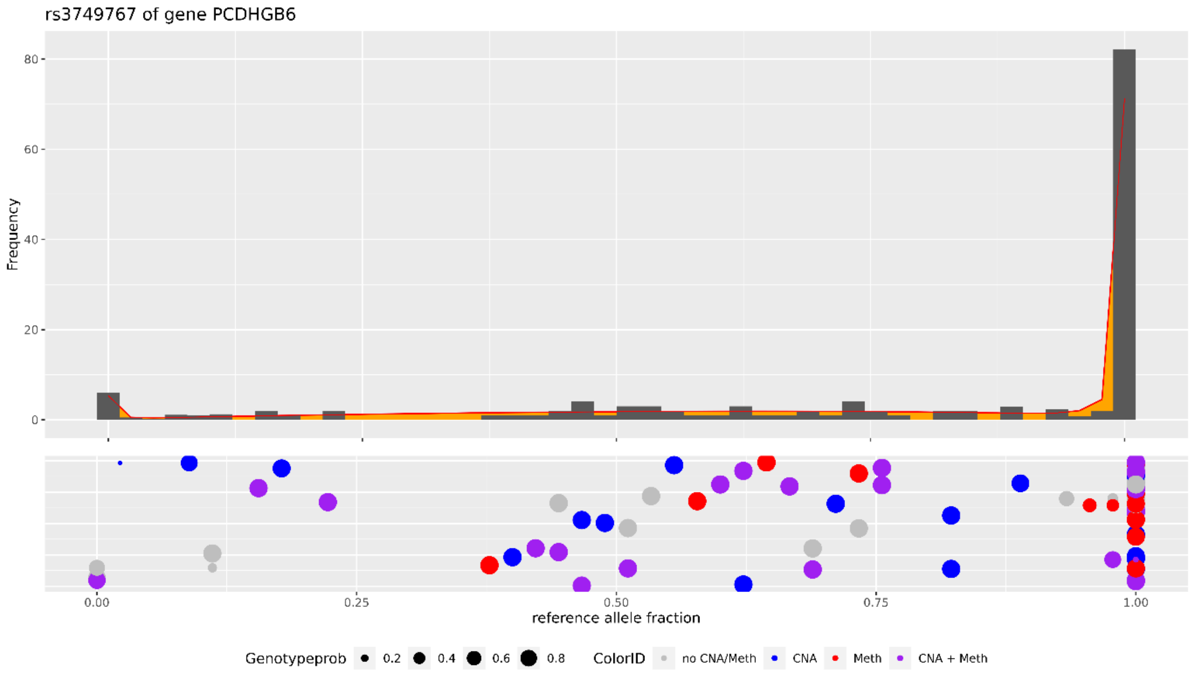
rs3749767 of gene PCDHGB6
Frequency
80
60
40
20
0
0.00
0.25
0.50
0.75
1.00
reference allele fraction
Genotypeprob
0.2
0.4
0.6
0.8
ColorID
no CNA/Meth
CNA
Meth
CNA + Meth

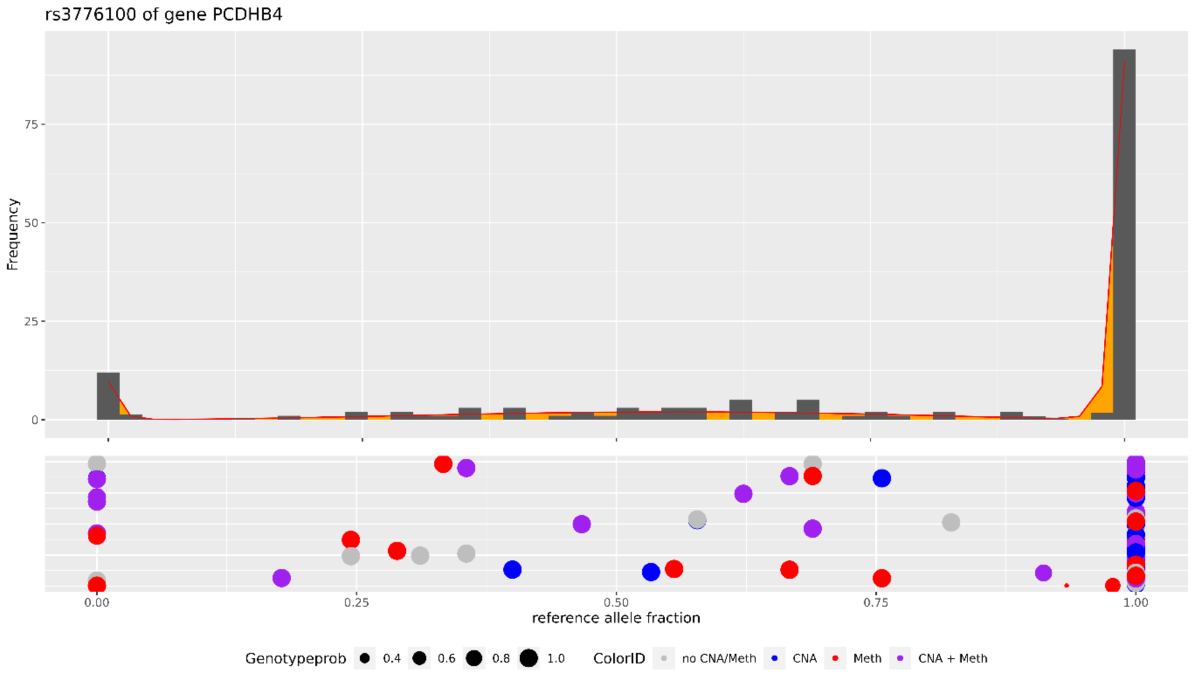
rs3776100 of gene PCDHB4
Frequency
75
50
25
0
0.00
0.25
0.50
0.75
1.00
reference allele fraction
Genotypeprob
0.4
0.6
0.8
1.0
ColorID
no CNA/Meth
CNA
Meth
CNA + Meth

##### X.3. Overdispersion estimation via joint- and separate model fit

For dAD-detection, MAGE fits both control- and case-overdispersion ($\rho$) via a joint model fit (i.e. other distributional parameters are shared between cases and controls; Section 4.1.C). Such a joint fit is necessary to construct a valid LRT for statistical dAD-detection, but MAGE is able to estimate overdispersion parameters on both the control- and case-populations separately as well, which can come in handy in e.g. RME studies. However, if the assumption of shared distributional parameters made during dAD-detection holds true, these fits' overdispersion estimates shouldn't differ much from one another, at least not to an extent that it would yield contradicting conclusions about the overdispersion of controls compared to cases.

Thus, as a check, we re-constructed the overdispersion plots in Figure 3 (Section 2.2.B) using $\rho$ values estimated from allele count fits on both the control- and case-population separately (summarized to gene-level in the same way as $\rho$ values from the regular fit; Section 4.2.F). The Figures below visualize this comparison for chromosome 3, showing the original joint fit (A; cf. Figure 3 and Section X.1) as well as the $\rho$ values from the separate fits (B). Unlike Figure 3, these are not color-coded according to statistical significance due to a LRT testing for significant dAD being impossible using the separate model fits.

The $\rho$ values from the separate model fits are generally lower than those from the joint model fit. This is to be expected, as fits of the remaining distributional parameters to the separate populations will of course fit those populations better, resulting in less surplus variance to be captured by the overdispersion parameter. The general trend of, and conclusions about, overdispersion in chromosome 3 (and all other chromosomes), however, remains the same.

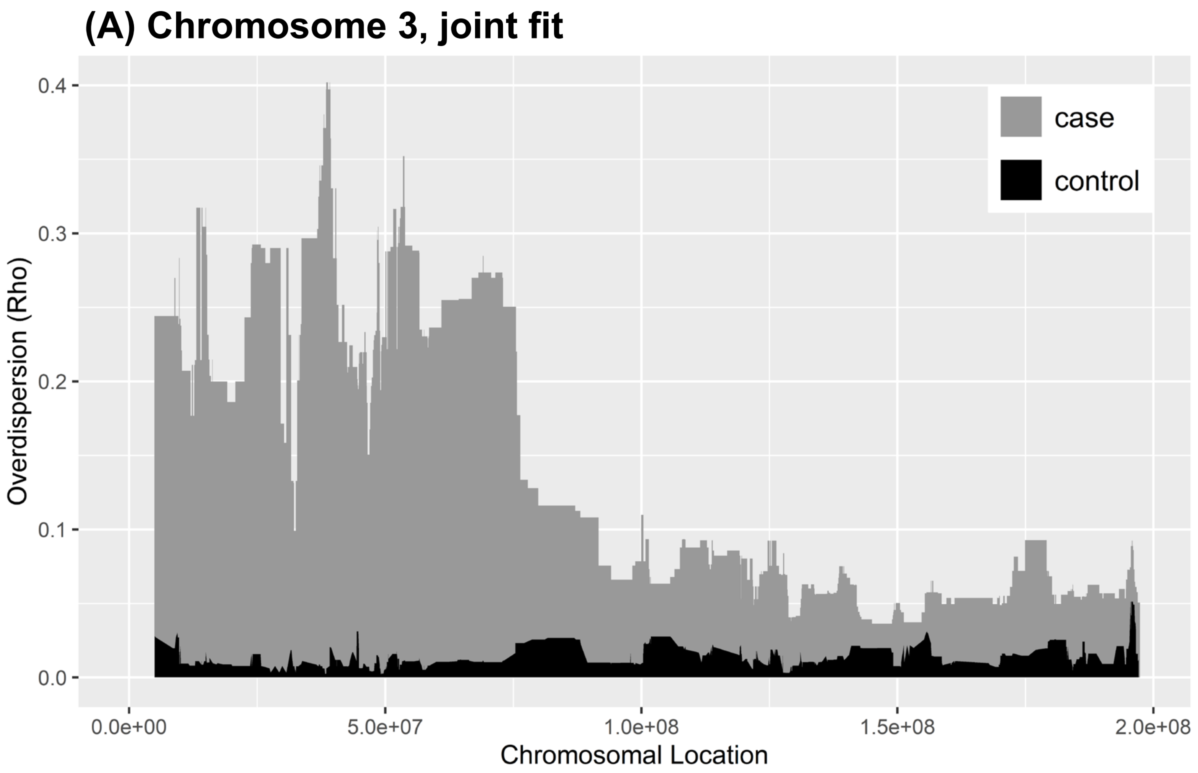

**(B) Chromosome 3, separate fit**

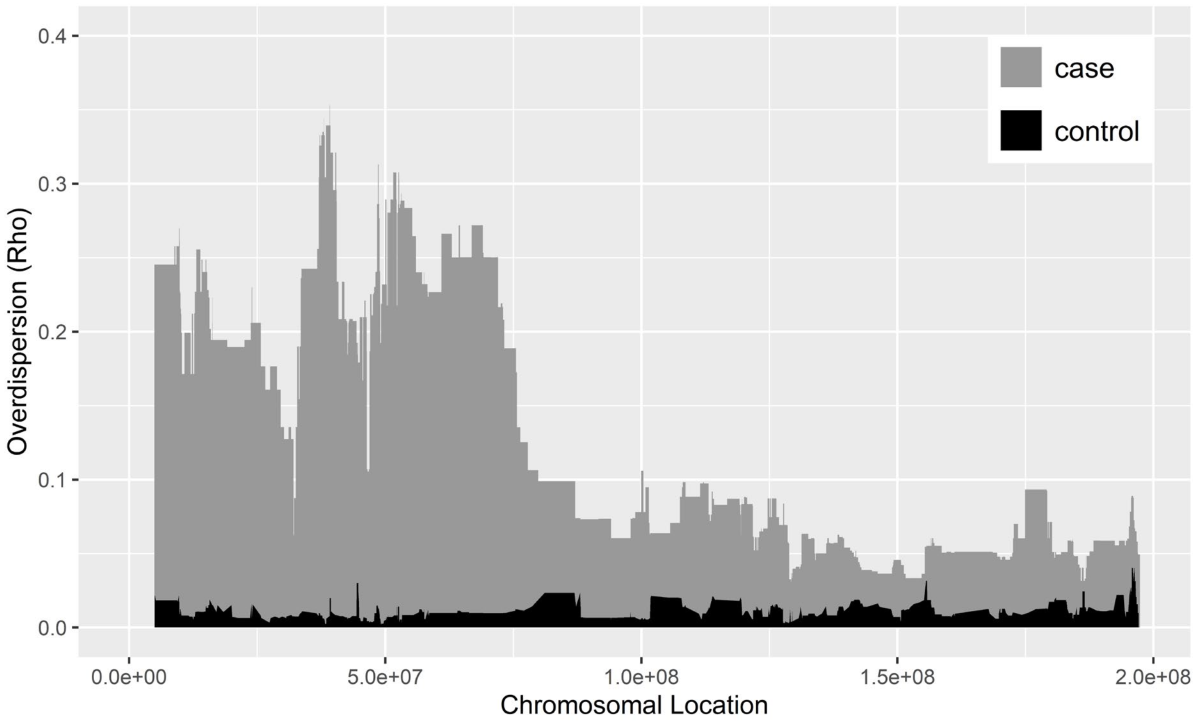
